# Engagement of the contralateral limb can enhance the facilitation of motor output by loud acoustic stimuli

**DOI:** 10.1101/2021.04.26.441558

**Authors:** Aaron N. McInnes, An T. Nguyen, Timothy J. Carroll, Ottmar V. Lipp, Welber Marinovic

## Abstract

When intense sound is presented during light muscle contraction, inhibition of the corticospinal tract is observed. During action preparation, this effect is reversed, with sound resulting in excitation of the corticospinal tract. We investigated how the combined maintenance of a muscle contraction during preparation for a ballistic action impacts the magnitude of the facilitation of motor output by a loud acoustic stimulus (LAS) – a phenomenon known as the StartReact effect. Participants executed ballistic wrist flexion movements and a LAS was presented simultaneously with the imperative signal in a subset of trials. We examined whether the force level or muscle used to maintain a contraction during preparation for the ballistic response impacted reaction time and/or the force of movements triggered by the LAS. These contractions were sustained either ipsilaterally or contralaterally to the ballistic response. The magnitude of facilitation by the LAS was greatest when low force flexion contractions were maintained in the limb contralateral to the ballistic response during preparation. There was little change in facilitation when contractions recruited the contralateral extensor muscle, or when they were sustained in the same limb that executed the ballistic response. We conclude that a larger network of neurons which may be engaged by a contralateral sustained contraction prior to initiation may be recruited by the LAS, further contributing to the motor output of the response. These findings may be particularly applicable in stroke rehabilitation where engagement of the contralesional side may increase the benefits of a LAS to the functional recovery of movement.

## 1.0 Introduction

The presentation of a loud acoustic stimulus (LAS) during movement preparation can affect the time of movement initiation as well as movement vigour. Actions that are sufficiently prepared at the time a LAS is delivered are involuntarily triggered at much shorter latencies, and are executed with greater force and vigour, than is typically produced voluntarily (Anzak, Tan, Pogosyan, & Brown, 2011; Anzak, Tan, Pogosyan, Djamshidian, et al., 2011; Marinovic et al., 2015; McInnes, Corti, et al., 2020; Valls-Solé et al., 1999). This is referred to as the StartReact effect (Valls-Solé et al., 1999). However, the effects of a LAS on motor circuits are contingent on the state of preparation for action at the time the stimulus is presented. For example, when the task is simply to maintain a light muscle contraction at a stable level, loud acoustic stimuli suppress the excitability of corticospinal pathways (Fisher et al., 2004; Furubayashi et al., 2000; Kuhn et al., 2004). In contrast, during a state of imminent preparation for a discrete action (i.e. the context in which the StartReact effect occurs), corticospinal excitability is increased shortly after the presentation of a LAS, which may provide a neurophysiological means by which motor output can be facilitated in the StartReact effect (Marinovic, Tresilian, et al., 2014). These observations highlight that the effects of a LAS on motor pathways are not fixed, but depend on the state of the motor system. However, the modulation of corticospinal excitability is further nuanced in that inhibition after acoustic stimulation is only observed when there is weak background muscle activity. During maintenance of a slightly stronger contraction, at 10% of maximum voluntary contraction (MVC), suppression of the corticospinal tract is less evident (Chen et al., 2016). This may be due to voluntary activation of the primary motor cortex (M1) suppressing intracortical inhibitory circuits as the amount of contraction force is increased (Roshan et al., 2003).

Furthermore, it is unclear how the maintenance of a muscle contraction during preparation for action may impact the StartReact effect. The potential observations which may be made under these conditions are currently uncertain as the effects of acoustic stimulation on the corticospinal tract during light muscle contraction are opposite (Fisher et al., 2004; Furubayashi et al., 2000; Kuhn et al., 2004) to those observed during action preparation (Marinovic, Tresilian, et al., 2014). Here, we investigated how different types of muscle contractions held during a preparatory foreperiod may impact the early triggering of motor actions and the enhancement of response vigour when the motor response is triggered by a LAS. If the combined maintenance of a muscle contraction during preparation for a subsequent action results in a decreased StartReact effect (i.e. reduced shortening of RT, reduced enhancement of response force/vigour), this would suggest that the contraction induces a suppressive effect of acoustic stimulation on motor pathways. In accordance with observations that the inhibitory LAS effect depends on the amount of background muscle activity, any putative reduction of the StartReact effect would be expected to be greatest at low contraction force levels. Alternatively, stable background contractions may increase preparatory neural activity prior to the discrete action and subsequently magnify the StartReact effect. During unilateral muscle contraction, excitability of the M1 ipsilateral to the contracting muscle increases as the amount of force is increased (Chen et al., 2019; Perez & Cohen, 2008; Shibuya et al., 2014; Stinear et al., 2001; Uematsu et al., 2010). In addition, regional cerebral blood flow in ipsilateral M1 decreases at light muscle contractions (5% of MVC) and increases in proportion to the strength of the muscle contraction from 10% - 60% of MVC (Dettmers et al., 1996). Therefore, contraction of a muscle during preparation of a contralateral response may result in an enhancement of the StartReact effect that is proportional to the strength of the contraction maintained during preparation. The StartReact effect has also been proposed as a tool to aid in rehabilitation in neurological conditions such as stroke, with startling sensory stimuli capable of reducing movement initiation-related deficits (Choudhury et al., 2019; Coppens et al., 2018; Honeycutt et al., 2015; Honeycutt & Perreault, 2012; Jankelowitz & Colebatch, 2004; Marinovic et al., 2016; Rahimi & Honeycutt, 2020). Given this, and the fact that stroke survivors typically experience exaggerated movement impairment on one side of the body (Zemke et al., 2003), maintenance of a contraction contralateral to the impaired side may be particularly beneficial for people with stroke if it enhances the benefits derived from intense sensory stimuli. Therefore, we examined how the type of isometric contraction maintained during preparation for a ballistic response impacts the facilitation of movement initiation and execution by a LAS, in both bilateral and unilateral tasks.

## 2.0 Method – Experiment one

### 2.1 Participants

Thirty participants were recruited for experiment one (20 female, mean age = 20.33 years, SD = 2.25). Participants in all experiments were self-reportedly right-handed, with normal or corrected-to-normal vision, and no apparent or known auditory impairments, neurological conditions, or injuries which may have affected their performance in the experiment. The study was approved by Curtin University’s local human research ethics committee and all participants provided informed, written consent before starting the experiment.

### 2.2 Procedures

Participants were seated on a height-adjustable chair with each hand and forearm secured in custom-made manipulanda, each housing a six degree of freedom force/torque sensor (JR3 45E15A-I63-A 400N60S, Woodland, CA; see de Rugy et al. (2012)). The forearm was secured in a semi-supinated position with the palms facing inward, and elbows flexed at an approximately 90° angle. Both hands and forearms of each participant were secured snugly in the device to prevent time delays between muscle activation and the recording of force. Participants were seated at a distance of approximately 0.8 m in front of a 24.5-inch monitor (Asus ROG Swift PG258Q, 120 Hz refresh rate, 1920×1080 resolution) which presented visual stimuli during the task. Both visual and auditory stimuli were presented using Psychtoolbox (v3.0.11) running in MATLAB 2015b.

Prior to the experimental trials, each participant completed a MVC procedure of wrist flexion in both the left and right hand (see Selvanayagam et al., 2016). In this procedure, force feedback was provided to subjects via a cursor in two-dimensional space (x = flexion/extension, y = abduction/adduction) such that 10 Newtons (N) was required to move the cursor 32 pixels on the computer monitor. Subjects produced three isometric MVCs for three seconds toward a target corresponding to the direction of wrist flexion, and the peak force was measured. The mean peak force of the three contractions for each hand was recorded as the MVC for the relevant hand. These data were used to determine the level of force required to reach targets during the experiment.

The experimental task required participants to perform a discrete ballistic wrist flexion movement of the right hand in response to an imperative cue. There were four contraction conditions during the experiment and these were each randomised across participants to one of four blocks of trials during the experiment. The contraction condition of the block determined the amount of force which was required to be sustained with the left hand during preparation of the right hand response. These force levels were 0%, 5%, 10%, and 20% of the participant’s left wrist flexor’s MVC. In one block, referred to as the “no contraction” condition, participants kept their left hand relaxed while they prepared and executed a ballistic flexion movement of the right hand, aiming to produce a brief force pulse of 20% of the right wrist flexor’s MVC. The 20% of MVC flexion ballistic response was chosen as we have previously shown that this muscle and force level is particularly prone to the beneficial effects of a LAS on motor output (McInnes, Corti, et al., 2020). In the three remaining contraction conditions, the left hand maintained an isometric flexion contraction at either 5%, 10%, or 20% of the left wrist flexor’s MVC, during preparation for the ballistic right-hand response. See Figure 1 for the sequence of events during the experiment. Prior to experimental trials, participants completed a block of 12 practice trials, which consisted of three trials of each condition in the experiment. Participants were given verbal feedback regarding their performance, and practice trials were repeated until participants were able to accurately initiate movements within 250 ms after the presentation of the imperative stimulus (IS). One-hundred and sixty-four experimental trials were then completed, split into four blocks of 41 trials each.

**Figure 1.**
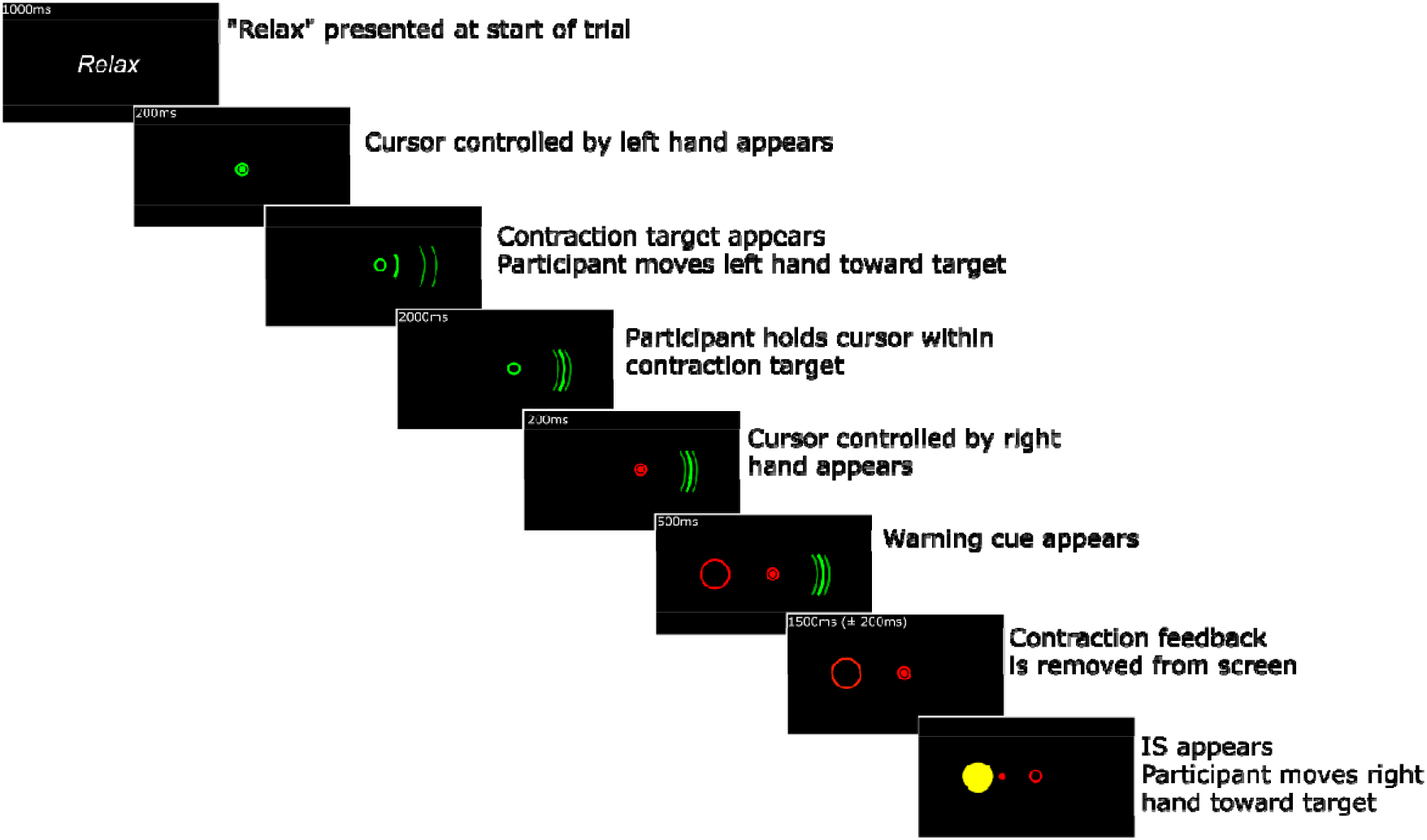
Sequence of events during the experiment requiring a left-hand contraction during preparation for a ballistic right hand response to the IS.

Each trial began with the word “relax” presented on-screen, indicating for the participant to keep their hands relaxed and stationary for the start of the trial. Next, a cursor that responded to forces with the left hand, and a contraction target, consisting of two arcs on the right side of the screen were presented. Participants moved their left hand so that the cursor was positioned within the contraction target, and held their hand in this position for the duration of the trial. The amount of force required to reach the contraction target changed each block, depending on the contraction condition (0%, 5%, 10%, or 20% of MVC contraction level). Trials would not proceed until the participant had maintained a contraction of the appropriate force level within a tolerance of ±7.5% of the target, to accommodate minor deviations from the contraction force. Once the participant had maintained this contraction for two seconds, a cursor which could be controlled by the right hand and a warning cue appeared, indicating the impending presentation of the IS. This warning cue appeared as a red circle on the left side of the screen. Participants were instructed to prepare to respond with the right hand during this period. After 500 ms, the contraction target and cursor indicating the position of the left hand was removed from the screen, so that participants would be encouraged to direct their attention to the warning cue and impending IS and prepare responses appropriately. If the participant unknowingly moved their left hand outside of the contraction target during this preparatory foreperiod, the left hand cursor and contraction target would reappear on screen, requiring the participant to return their left hand within the contraction target before the trial would proceed. The warning cue was presented for two seconds (± 200 ms jitter), after which the IS, a yellow circle in place of the warning cue, was presented. Twenty percent of trials occurred as probe trials in which a LAS was presented as an accessory stimulus simultaneously with the IS. The order of trials was pseudo-randomised so that the LAS would not be presented in two consecutive trials. When the IS was presented, participants reacted by moving their right hand in a ballistic wrist flexion. They were instructed to aim to touch the target and stop the cursor movement once the target had been reached. To encourage participants to respond with the appropriate amount of force required to reach the target, the yellow IS target flashed green when intersected with the cursor. At the end of the trial, feedback regarding RT was presented to encourage quick responses throughout the experiment. In probe trials, this feedback was not presented.

### 2.3 Loud Acoustic Stimulus

In probe trials, a LAS generated by the onboard audio of the computer used to run experiments was presented through high-fidelity stereophonic active noise cancelling headphones (Bose QC25). The peak amplitude of the stimulus was measured at 105 dBa using a Bruel and Kjaer sound level meter (Type 2205, A weighted; Brüel & Kjaer Sound and Vibration Measurement, Naerum, Denmark). The LAS was presented for a duration of 50 ms and with a rise and fall time < 1.5 ms.

### 2.4 Statistical analyses

For each trial, the time series of force data were collected from the load cell with a sampling rate of 2 kHz using a National Instruments data acquisition device (NI USB-6229). Movement onsets were estimated from the force time series data using Teasdale et al.’s (1993) algorithm, and RT was determined by subtracting the time of IS presentation from the time of movement onset. The vigour of the ballistic response was determined by measuring the derivative of the torque data with respect to time, referred to as the rate of force development (Newtons per second; N/s). Peak rate of force development and peak force were determined as the maximum values of the force/force derivative time-series data reached over the course of a trial. Statistical analyses were run using R software (*v3.6.0*; R Core Team, 2019).

Prior to analysis, trials with a RT < 60 ms or > 1000 ms were removed on the basis that these were error responses made as a result of premature response initiation due to anticipation of the IS, or delayed responses due to insufficient movement preparation (Whelan, 2008). This resulted in the exclusion of 100 trials (2.03% of all trials) in experiment one. We further used cumulative distribution functions (CDFs) to analyse mean RTs at each percentile of the entire RT distribution to assess whether preparatory contraction conditions resulted in movements being more or less prone to triggering delays. We have outlined the method of analysing data using CDFs in a StartReact context in more detail previously (McInnes, Castellote, et al., 2020).

We used the *lmer* function from the *lmerTest* package (*v3.1*; Kuznetsova, Brockhoff, & Christensen, 2017) to conduct a series of linear mixed-effects models. All trials were fed into the linear models with participants set as a random factor. In Experiment one, trial type (control, LAS) and contraction level (no contraction, 5%, 10%, 20% contractions) were fixed factors in the model. To determine the extent of facilitation for RT, peak force, and peak rate of force development that occurred as a result of the LAS, we calculated differences in RT and ratios of peak force and peak rate of force development. We analysed both raw values and differences/ratios as raw values provided a demonstration of the potential sustained contraction effects on voluntary responses in control trials, while differences/ratios illustrated how the magnitude of the StartReact effect may be impacted by the contractions sustained during preparation. For RT differences, the median RT of control trials was calculated for each contraction condition, and each LAS trial was subtracted from the median of control trials to determine a RT difference for each probe trial. A similar procedure was conducted with peak force and peak rate of force development by dividing probe trials by the median of control trials. For all models, degrees of freedom were approximated using the Kenward-Roger procedure. *R*^2^ values, calculated using the *r2beta* function (r2glmm package, v0.1.2) are also reported to estimate effect sizes of all main effects and interactions tested using the linear mixed models. Post-hoc tests were conducted for significant main effects and interactions of the linear mixed models using the *emmeans* function (emmeans package, v1.3.0) with the correction of multiple comparisons using the false discovery rate method (Benjamini & Hochberg, 1995).

## 3.0 Results – Experiment 1

### 3.1 Shortening of response initiation

RT was significantly shortened in LAS probe trials (*M* = 158.69 ms, *SD* = 85.37) in comparison to control trials (*M* = 221.74 ms, *SD* = 99.86), with a statistically significant main effect of trial type for RT, *F*_(1, 4783.1)_ = 639.84, *p* < .001, *R*^2^ = .118. The main effect of contraction level was also statistically significant for RT, *F*_(3, 4783.2)_ = 9.50, *p* < .001, *R*^2^ = .006. The interaction of trial type with contraction level was not statistically significant, *F*_(3,_ 4783.1) = 0.22, *p* = .886, *R*^2^ < .001. Analysis of the difference in RT between probe trials and the median of control trials for each condition did not indicate a statistically significant main effect of contraction level, *F*_(3, 887.49)_ = 0.65, *p* = .583, *R*^2^ = .002. Mean RTs across each condition are shown in Figure 2.

**Figure 2.**
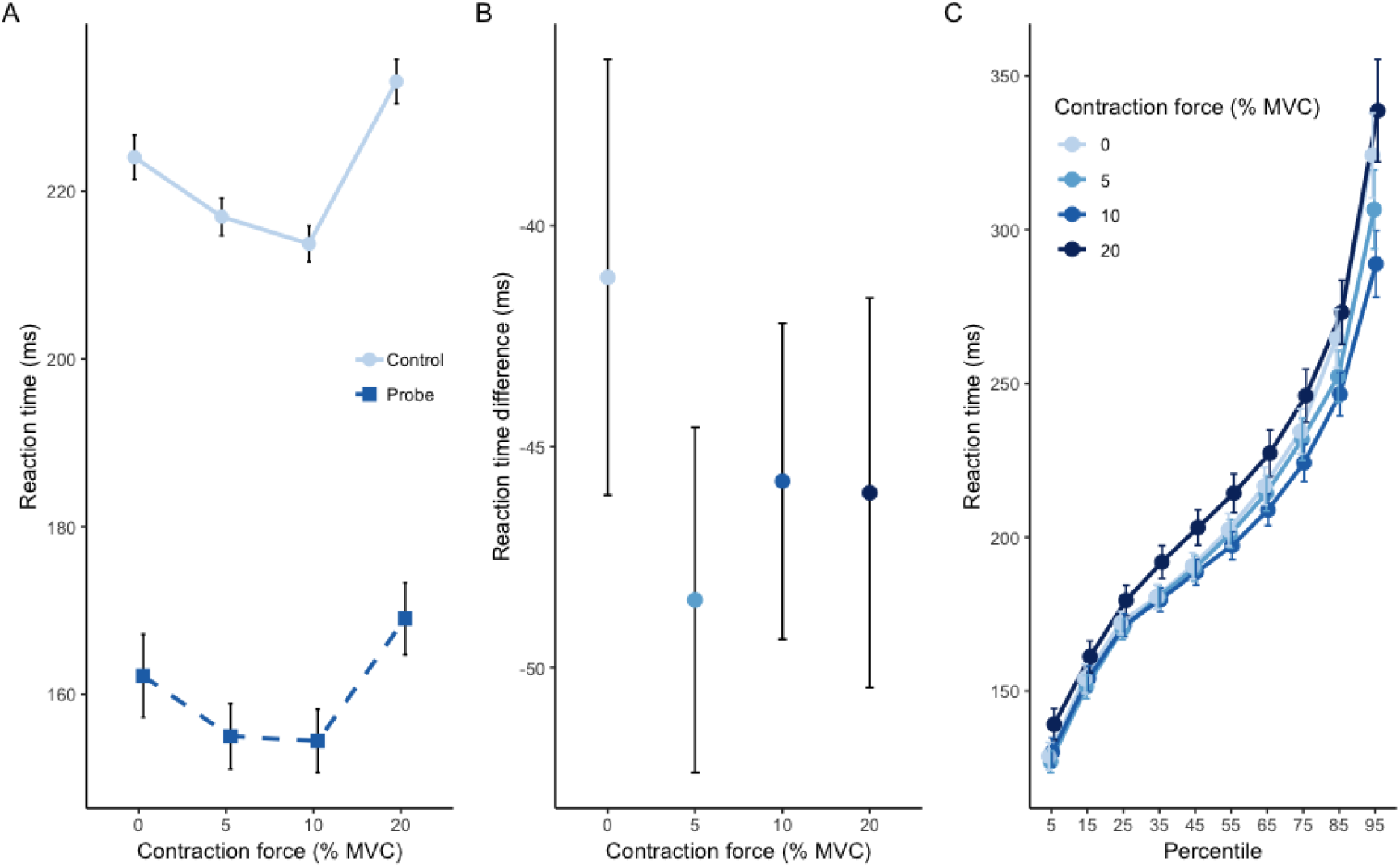
A). Mean reaction time over control and probe trials for each contraction level. B). Mean of the difference in RT between all probe trials and the median of control trials for each condition. C). Mean RT across each participant for each percentile of RT. Error bars represent standard error of the mean.

We further examined each participant’s mean RT at each percentile, across the four contraction level conditions. A linear mixed-effects model indicated a significant main effect of contraction level, *F*_(3, 1131)_ = 24.10, *p* < .001, *R*^2^ = .060. In comparison to the no contraction condition (*M* = 206.21 ms, *SD* = 70.40), the 10% contraction condition showed significantly shorter RTs across the CDF curve (*M* = 198.76 ms, *SD* = 57.25; *p* = .003), while the 20% contraction condition had significantly longer RTs (*M* = 217.18 ms, *SD* = 73.51; *p* < .001). RTs across the CDF curve for the 5% contraction condition (*M* = 201.87 ms, *SD* = 65.48) were not significantly different from the no contraction condition (*p* = .083), nor the 10% contraction condition (*p* = .180). The interaction of percentile with contraction level was not statistically significant, *F*_(27, 1131)_ = 1.28, *p* = .153, *R*^2^ = .030.

### 3.2 Enhancement of peak force and vigour

Peak force showed an enhancement in probe trials (*M* = 36 N, *SD* = 16.60) compared to control trials (*M* = 31.95 N, *SD* = 13.31), as shown by the main effect of trial type which was statistically significant, *F*_(1, 4783)_ = 127.09, *p* < .001, *R*^2^ = .026. The main effect of contraction level was also statistically significant, *F*_(3, 4783)_ = 10.59, *p* < .001, *R*^2^ = .007. Furthermore, the interaction of trial type and contraction level was statistically significant for peak force, *F*_(3,_ 4783) = 6.83, *p* < .001, *R*^2^ = .004. Analysis of the ratios of peak force showed a statistically significant main effect of contraction level, *F*_(3, 908.26)_ = 6.35, *p* < .001, *R*^2^ = .021. Post hoc tests indicated that in comparison to the no contraction condition (*M* = 1.11, *SD* = 0.39), ratios of peak force between control trials and probe trials were significantly greater in the 10% contraction condition (*M* = 1.19, *SD* = 0.33; *p* = .016), but not in the 5% (*M* = 1.17, *SD* = 0.40; *p* = .059) or 20% contraction conditions (*M* = 1.07, *SD* = 0.36; *p* = .417).

Similarly to peak force, our analysis showed a statistically significant main effect of trial type for peak rate of force development, *F*_(1, 4783)_ = 252.01, *p* < .001, *R*^2^ = .050, with greater peak rate of force development observed on average for LAS probe trials (*M* = 492.13 N/s, *SD* = 251.70) in comparison to control trials (*M* = 410.50 N/s, *SD* = 185.05). The main effect of contraction level, *F*_(3, 4783)_ = 4.56, *p* = .003, *R*^2^ = .003, and the interaction of trial type with contraction level, *F*_(3, 4783)_ = 5.43, *p* = .001, *R*^2^ = .003, were statistically significant. The main effect of contraction level for ratios of peak rate of force development was also statistically significant, *F*_(3, 908.25)_ = 5.46, *p* = .001, *R*^2^ = .018. In comparison to the no contraction condition (*M* = 1.17, *SD* = 0.52), post hoc tests indicated ratios of peak rate of force development were significantly greater for the 5% (*M* = 1.25, *SD* = 0.44; *p* = .042) and 10% (*M* = 1.29, *SD* = 0.42; *p* = .006) contraction conditions but not for the 20% contraction condition (*M* = 1.16, *SD* = 0.45; *p* = .817). The mean peak force and vigour for each experimental condition are presented in Figure 3.

**Figure 3.**
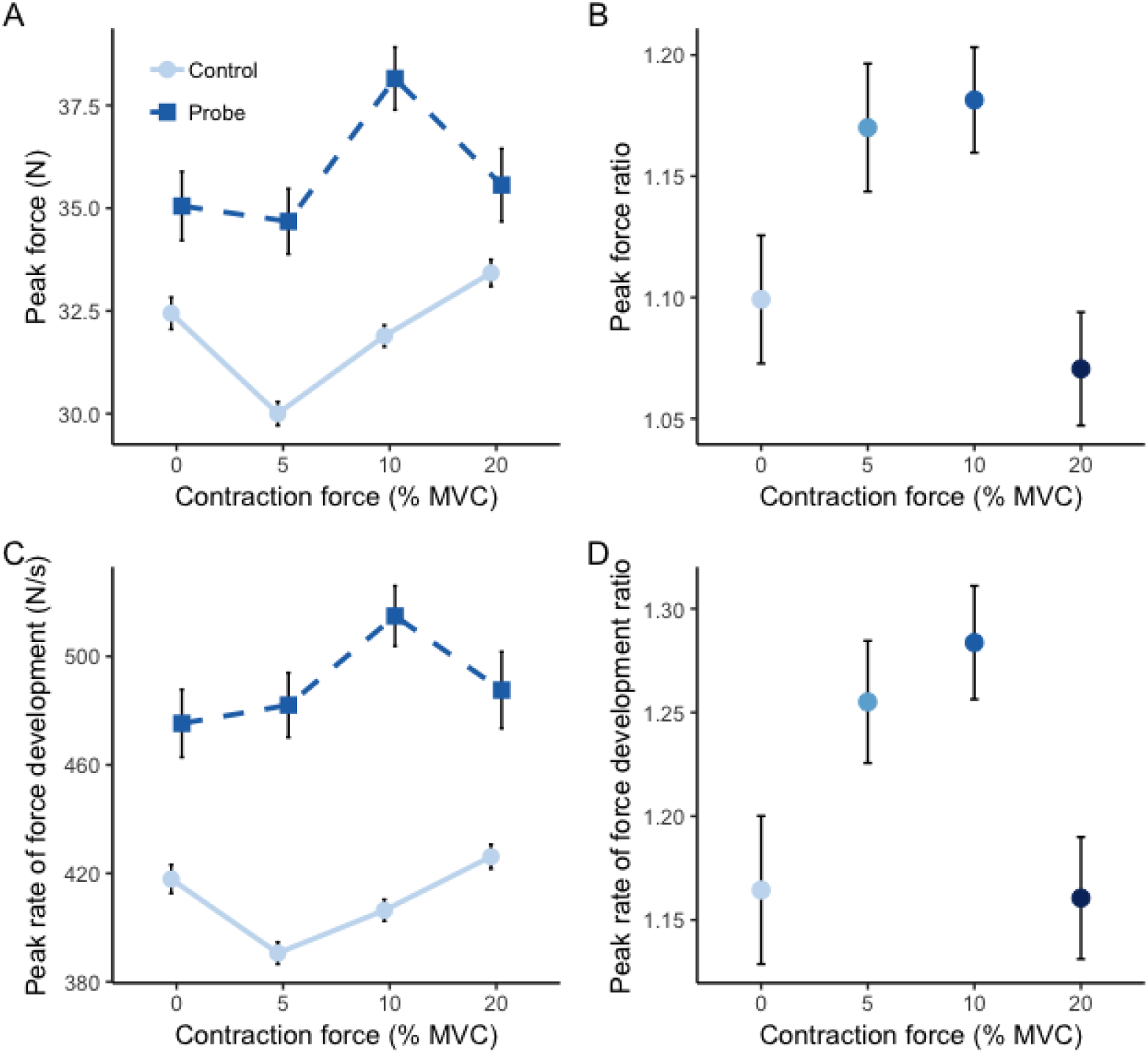
A). Mean peak force for control and probe trials over each contraction level. B). Mean peak force ratios over each contraction level. C). Mean peak rate of force development over control and probe trials at each contraction level. D). Mean peak rate of force development ratios for each contraction level. Error bars represent standard error of the mean.

## 6.0 Method – Experiment two

### 6.1 Participants

A second sample of 25 participants (16 female; mean age = 20.28 years, SD = 1.65) was recruited for experiment two. The same recruitment criteria as experiment one was used for experiment two.

### 6.2 Procedures

In experiment two, we used a forewarned RT task similar to experiment one, in a unilateral task. In this task, isometric contractions were maintained during preparation for the response to the IS with the same (right) hand. Responses to the IS were again ballistic flexion movements of the right hand at 20% of MVC of the right wrist flexor. The right (responding) hand either remained relaxed during preparation for the response to the IS, or maintained a contraction in either flexion or extension, at 10% of the relevant muscle’s MVC. Both flexion and extension contractions were examined in order to assess whether the potential observable effects were muscle specific. Flexion and extension contractions were employed as these are an agonist/antagonist pair and have also been suggested to differ in the strength of their efferent contributions from the corticospinal and reticulospinal tracts (Cheney & Fetz, 1980; Clough et al., 1968; de Noordhout et al., 1999; Fetz & Cheney, 1980; Godfrey et al., 2013; Koganemaru et al., 2010; McInnes, Castellote, et al., 2020; McMillan et al., 2004; Palmer & Ashby, 1992; Park & Li, 2013; Quinn et al., 2018; Vallence et al., 2012). Contractions were maintained during preparation at 10% of the muscle’s MVC as this force level appeared to provide the most benefit in experiment one. The ballistic response always required additional responsive activation of the flexor muscle at 20% of MVC beyond the contraction position. For example, during the isometric flexion contraction, the target was set so that from the 10% of MVC contraction position, an additional force of 20% of flexion MVC would be required to meet the target (i.e. the final position of the ballistic response was 30% of flexion MVC). During the isometric extension contraction, the ballistic flexion response of 20% of MVC was required, measured from the point at which the extensor muscle was at rest (i.e. the final force target required force away from 10% extension MVC and toward 20% flexion MVC). We determined this to be the most feasibly equivalent between the flexion and extension contraction conditions of the unilateral task in terms of the amount of force beyond the sustained contraction force which would be required to generate the final ballistic response. Trials were again excluded from analysis on the basis of 60 ms < RT > 1000 ms. This resulted in the exclusion of 98 trials from experiment two in total (1.96% of all trials). In addition, we calculated the variability of force 250 ms prior to the presentation of the IS by calculating the standard deviation of force at this time point. This analysis was conducted for this experiment as differences in the variability of force during the sustained contraction prior to the ballistic force may have impacted detection of force onset for the ballistic response during the unilateral task. As a result, this potential systematic influence of movement onset detection may have impacted our analysis of RT in this experiment. Therefore, we fed force variability into a linear mixed model to examine whether force variability prior to the IS systematically differed as a function of the contraction condition.

## 7.0 Results - Experiment two

### 7.1 Shortening of response initiation

RTs were significantly shorter in probe trials (*M* = 159.98 ms, *SD* = 101.61) in comparison to control trials (*M* = 238.25 ms, *SD* = 140.42), with a statistically significant main effect of trial type for RT, *F*_(1, 3648)_ = 406.13, *p* < .001, *R*^2^ = .100. The main effect of contraction type was also statistically significant for RT, *F*_(2, 3648.1)_ = 10.97, *p* < .001, *R*^2^ = .006. Responses on average showed shorter RTs in the no contraction condition (*M* = 209.65 ms, *SD* = 107.1) in comparison to when contractions were maintained in both flexion (*M* = 223.97 ms, *SD* = 123.41; *p* = .012) and extension (*M* = 233.58 ms, *SD* = 111.75; *p* < .001) during preparation. However, analysis of the variability of force 250 ms prior to the IS indicated a significant main effect of contraction type, *F*_(2, 3648)_ = 34.88, *p* < .001, *R*^2^ = .021. This indicates this effect of contraction type on RT may be an artefact of more variable baseline force affecting the detection of movement onset in this experiment. Force variability 250 ms prior to the IS was significantly lower for the no contraction condition (*M* = 0.04, *SD* = 0.25) in comparison to both the flexion (*M* = 0.09, *SD* = 0.14; *p* < .001) and extension contraction conditions (*M* = 0.08, *SD* = 0.09; *p* < .001). Contraction force variability was also significantly different between flexion and extension contractions (*p* = .032). The interaction of trial type with contraction type was not statistically significant, *F*_(2, 3648)_ = 0.21, *p* = .811, *R*^2^ = .002. Analysis of the difference in RT between probe trials and the median of control trials for each contraction condition did not indicate a statistically significant main effect of contraction type, *F*_(2, 711.16)_ = 1.23, *p* = .293, *R*^2^ = .003.

Each participant’s mean RT at each percentile contraction conditions was also analysed using a linear mixed-effects model. Analysis indicated a significant main effect of contraction type, *F*_(2, 696)_ = 25.38, *p* < .001, *R*^2^ = .068. In comparison to the no contraction condition (*M* = 204.08 ms, *SD* = 89.27), the flexion contraction condition showed significantly longer RTs across the CDF curve (*M* = 219.82 ms, *SD* = 113.3; *p* < .001), as did the extension contraction condition (*M* = 230.83 ms, *SD* = 107.94; *p* < .001). RTs across the CDF curve were significantly longer for the extension contraction condition in comparison to the flexion contraction condition (*p* = .003). The interaction of percentile with contraction type was not statistically significant, *F*_(18, 696)_ = 1.17, *p* = .282, *R*^2^ = .029. Figure 4 shows the mean RTs for each condition along with mean RTs at each percentile within the CDF.

**Figure 4.**
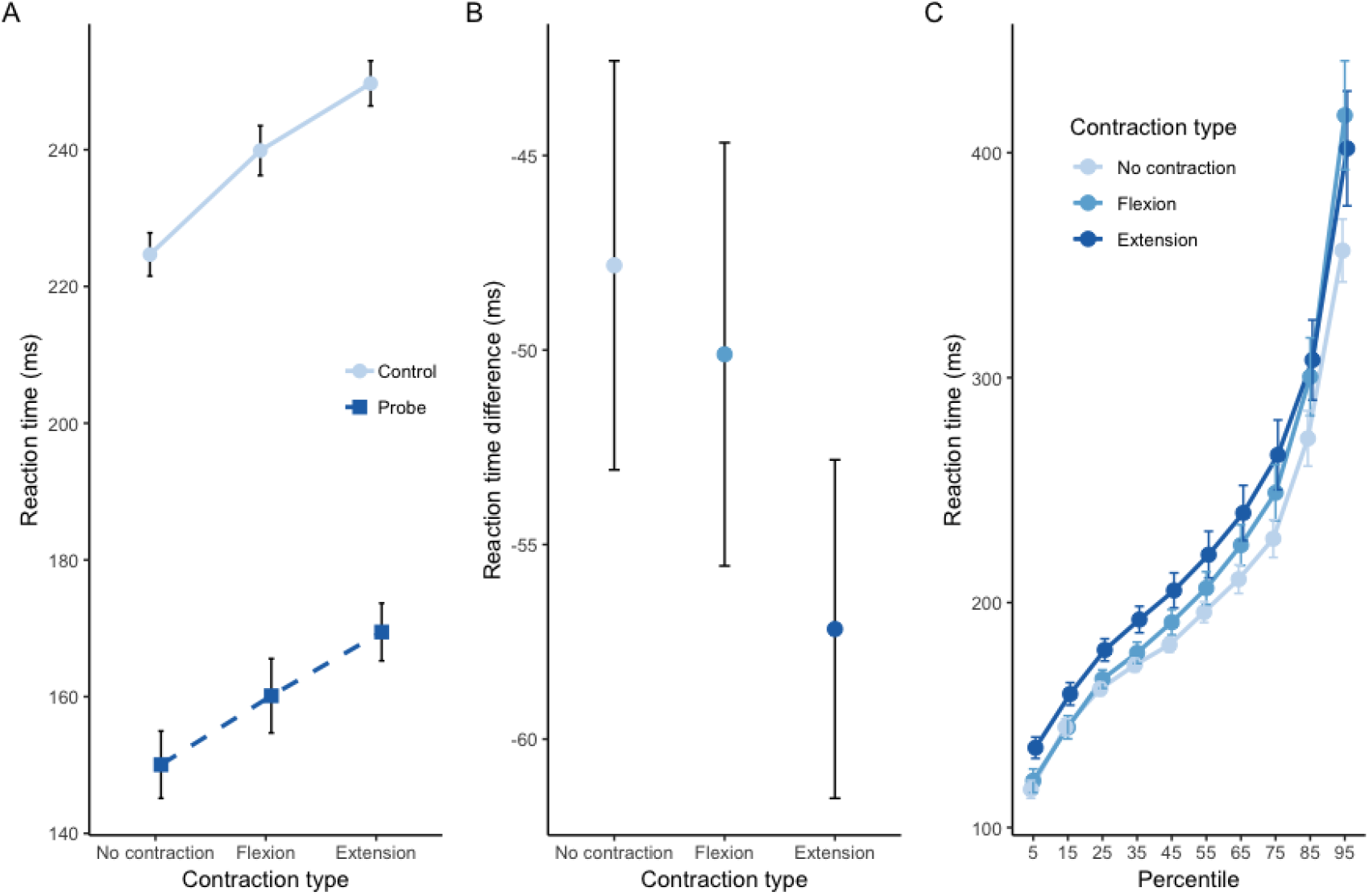
A). Mean reaction time over control and probe trials for each contraction type. B). Mean difference in RT between control and probe trials for each contraction type. C). B). Mean RT across each participant for each percentile of RT. Error bars represent standard error of the mean.

### 7.2 Facilitation of response force and vigour

Peak force showed an enhancement in probe trials (*M* = 44.34 N, *SD* = 18.33) compared to control trials (*M* = 39.24 N, *SD* = 16.52), as shown by a statistically significant main effect of trial type, *F*_(1, 3648)_ = 111.54, *p* < .001, *R*^2^ = .030. The main effect of contraction type was also statistically significant, *F*_(2, 3648)_ = 66.22, *p* < .001, *R*^2^ = .035. The flexion (*M* = 37.46 N, *SD* = 12.67) contraction condition showed significantly lower peak force on average in comparison to both the no contraction (*M* = 37.46 N, *SD* = 12.69; *p* < .001) and extension (*M* = 43.84 N, *SD* = 15.38; *p* < .001) contraction conditions. Average peak force in the extension contraction condition was also significantly greater than the no contraction condition (*p* < .001). The interaction of trial type with contraction type was not statistically significant, *F*_(2, 3648)_ = 0.23, *p* = .793, *R*^2^ = .000. The benefit of the acoustic stimulus on peak force did not appear to differ as a function of contraction type, as analysis of the ratios of peak force indicated the main effect of contraction type was not statistically significant, *F*_(2, 711.14)_ = 1.02, *p* = .361, *R*^2^ = .003.

Peak rate of force development was also increased by the LAS (*M* = 567.93 N/s, *SD* = 288.57) in comparison to control trials (*M* = 448.40 N/s, *SD* = 218.79), as indicated by a main effect of trial type, *F*_(1, 3648)_ = 313.27, *p* < .001, *R*^2^ = .079. The main effect of contraction type was also statistically significant, *F*_(2, 3648)_ = 39.12, *p* < .001, *R*^2^ = .021, however, the interaction of trial type with contraction type, *F*_(2, 3648)_ = 1.30, *p* = .273, *R*^2^ = .001, was not significant. The main effect of contraction type for ratios of peak rate of force development was statistically significant, *F*_(2, 711.15)_ = 5.83, *p* = .003, *R*^2^ = .016. Post hoc tests indicated peak rate of force development ratios in the flexion (*M* = 1.40, *SD* = 0.54; *p* = .002), but not the extension (*M* = 1.33, *SD* = 0.54; *p* = .088) contraction condition, were significantly greater than the no contraction condition (*M* = 1.26, *SD* = 0.49). The flexion contraction condition also showed significantly greater peak rate of force development ratios in comparison to the extension contraction condition (*p* = .088). Ratios and means of peak force and peak rate of force development are presented in Figure 5.

**Figure 5.**
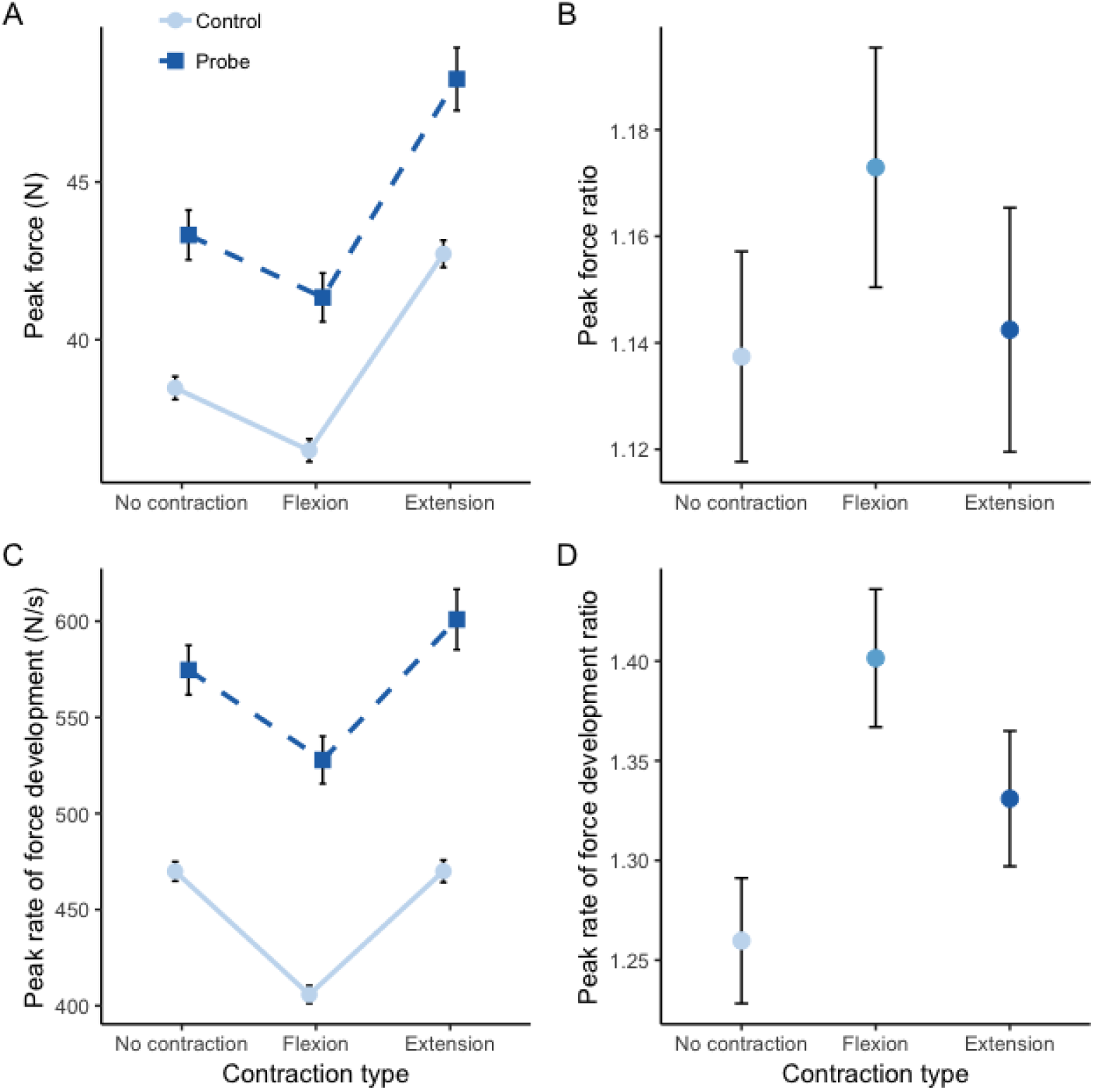
A). Mean peak force for control and probe trials for each contraction type. B). Mean peak force ratios for each contraction type. C). Mean peak rate of force development over control and probe trials at each contraction type. D). Mean peak rate of force development ratios for each contraction type. Error bars represent standard error of the mean.

## 4.0 Method – Experiment three

### 4.1 Participants

In experiment three, a sample of 29 volunteers (different from those recruited in experiments one and two) were recruited (23 female, mean age = 20.72 years, SD = 3.18). Participants were again required to be right-handed and free from any auditory impairments, neurological conditions, or injuries which may have impacted their performance in the experiment.

### 4.2 Procedures

Experiment three followed similar procedures to those of the bilateral task of experiment one, except contractions were made with the left hand in both directions of flexion and extension during preparation in an anticipatory timing task requiring a response from the right hand. The use of both flexion and extension contractions during preparation in the bimanual task again allowed us to examine whether the effects observed in experiment one were muscle specific. Alternatively, effects may be movement specific. For example, modulations of corticospinal excitability in M1 during ipsilateral movement have been suggested to be more strongly associated with whether the direction of movement is toward or away from the midline of the body, rather than the specific agonist muscle used in the movement (Duque et al., 2005). An anticipatory timing task was used in this experiment as the effects of the LAS on peak force and vigour were larger than those observed for the latency of movement onset in experiment one. Therefore, an anticipatory timing protocol allowed us to examine whether the effects of the LAS on movement execution become more or less pronounced when the stimulation is delivered closer in time to movement onset. We presented contraction feedback as an outer ring of a circle, with the contraction target at the 12 o’clock position of the circle. Rather than the presentation of a WS and IS, as in the previous experiments, the centre of the contraction feedback would fill in according to a clockwise motion, and participants were instructed to initiate their movement in synchrony with the time at which the circle was completely filled and the clock hand intersected at the 12 o’clock position. Contractions during preparation were set at one required force level – 10% of MVC, as this force level provided the most benefit in experiment one. As in the previous experiments, responses were made with the right hand at 20% of MVC. The LAS was presented in synchrony with the expected time of movement onset and, therefore, we did not anticipate a main effect of LAS on the temporal error of movement onset. In experiment three, responses to the IS with temporal error < −150 ms or > 150 ms were excluded from analysis. This resulted in the exclusion of 519 trials in experiment three (12.14% of all trials).

## 5.0 Results - Experiment three

### 5.1 Temporal error of movement onset

Mean temporal error of movement onset was earliest in the no contraction condition (*M* = −24.83 ms, *SD* = 51.13), followed by that of the extension contraction (*M* = −16.68 ms, *SD* = 50.50) and flexion contraction (*M* = −10.77 ms, *SD* = 50.18) conditions. Our analysis of temporal error of movement onset data in experiment three indicated a statistically significant main effect of contraction type (no contraction/flexion/extension), *F*_(2, 3733.4)_ = 22.76, *p* < .001, *R*^2^ = .012. The time of movement initiation in the no contraction condition was significantly earlier than both the flexion (*p* < .001) and extension (*p* < .001) contraction conditions. The difference in temporal error between the flexion and extension contraction conditions was also statistically significant (*p* = .006). As expected given the timing of LAS presentation, the main effect of trial type (LAS/control) was not statistically significant, *F*_(1,_ 3732.5) = 2.59, *p* = .107, *R*^2^ = .001, nor was the interaction of trial type with contraction type, *F*_(2, 3732.4)_ = 0.02, *p* = .977, *R*^2^ = .002. Mean temporal error for each condition is shown in Figure 6.

**Figure 6.**
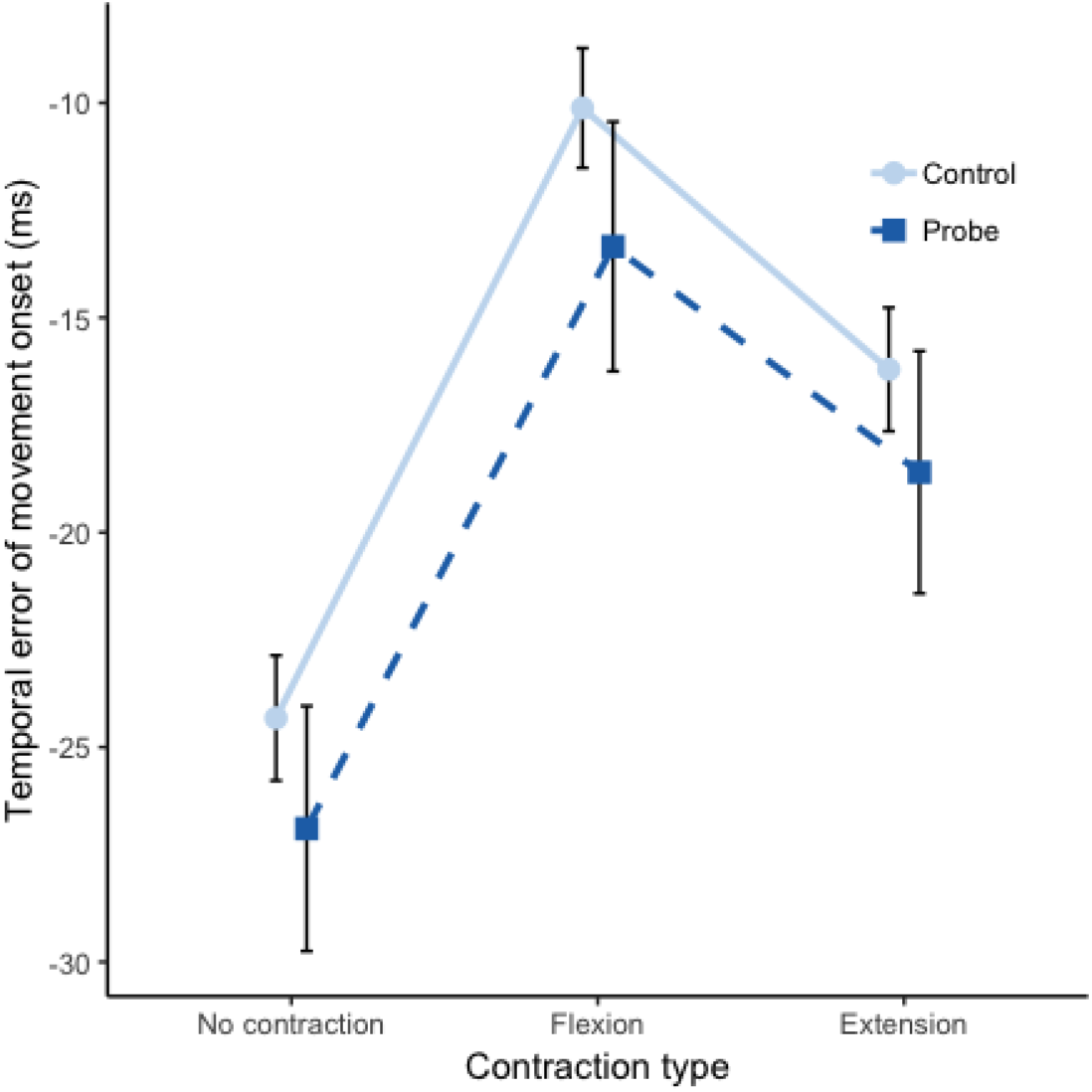
Mean temporal error of movement onset for control and probe trials across contraction conditions. Error bars represent standard error of the mean.

### 5.2 Enhancement of response force and vigour

Peak force was enhanced by the LAS (*M* = 27.9 N, *SD* = 14.03) in comparison to control trials (*M* = 23.54 N, *SD* = 9.64). The main effect of trial type for peak force was statistically significant, *F*_(1, 3732)_ = 197.77, *p* < .001, *R*^2^ = .050. The main effect of contraction type was not statistically significant, *F*_(2, 3732)_ = 2.38, *p* = .092, *R*^2^ = .001. Furthermore, the interaction of trial type with contraction type failed to reach statistical significance, *F*_(2, 3732)_ = 1.84, *p* = .159, *R*^2^ = .001. Ratios of peak force were largest in the flexion contraction condition (*M* = 1.26, *SD* = 0.51), with smaller ratios of peak force being found for the no contraction (*M* = 1.19, *SD* = 0.42) and extension contraction conditions (*M* = 1.18, *SD* = 0.47). A linear mixed model of peak force ratios indicated a significant main effect of contraction type, *F*_(2, 725.52)_ = 3.34, *p* = .036, *R*^2^ = .009. Post hoc tests indicated a significant difference in peak force ratios between the flexion contraction condition and the no contraction condition (*p* = .049), between the flexion contraction condition and the extension contraction condition (*p* = .049), but not between the no contraction and extension contraction conditions (*p* = .872).

On average, peak rate of force development was also increased by the LAS (*M* = 347.98 N/s, *SD* = 225.50) in comparison to control trials (*M* = 258.41 N/s, *SD* = 121.88). Linear mixed-effects models of peak rate of force development indicated a significant main effect of trial type, *F*_(1, 3732)_ = 435.27, *p* < .001, *R*^2^ = .104. The main effect of contraction type, *F*_(2, 3732)_ = 4.44, *p* = .012, *R*^2^ = .002, as well as the interaction of trial type with contraction type, *F*_(2, 3732)_ = 7.90, *p* < .001, *R*^2^ = .004, were statistically significant. Analysis of the ratios of probe trials over control trials for peak rate of force development indicated a significant main effect of contraction type, *F*_(2, 725.39)_ = 7.22, *p* < .001, *R*^2^ = .020, with larger ratios of peak rate of force development occurring in the flexion contraction (*M* = 1.49, *SD* = 0.80, *p* < .001) and extension contraction (*M* =1.41, *SD* = 0.72, *p* = .032) conditions in comparison to the no contraction condition (*M* = 1.29, *SD* = 0.63). The difference between the flexion contraction and extension contraction conditions was not statistically significant (*p* = .150). The means and ratios of peak force and peak rate of force development are shown in Figure 7.

**Figure 7.**
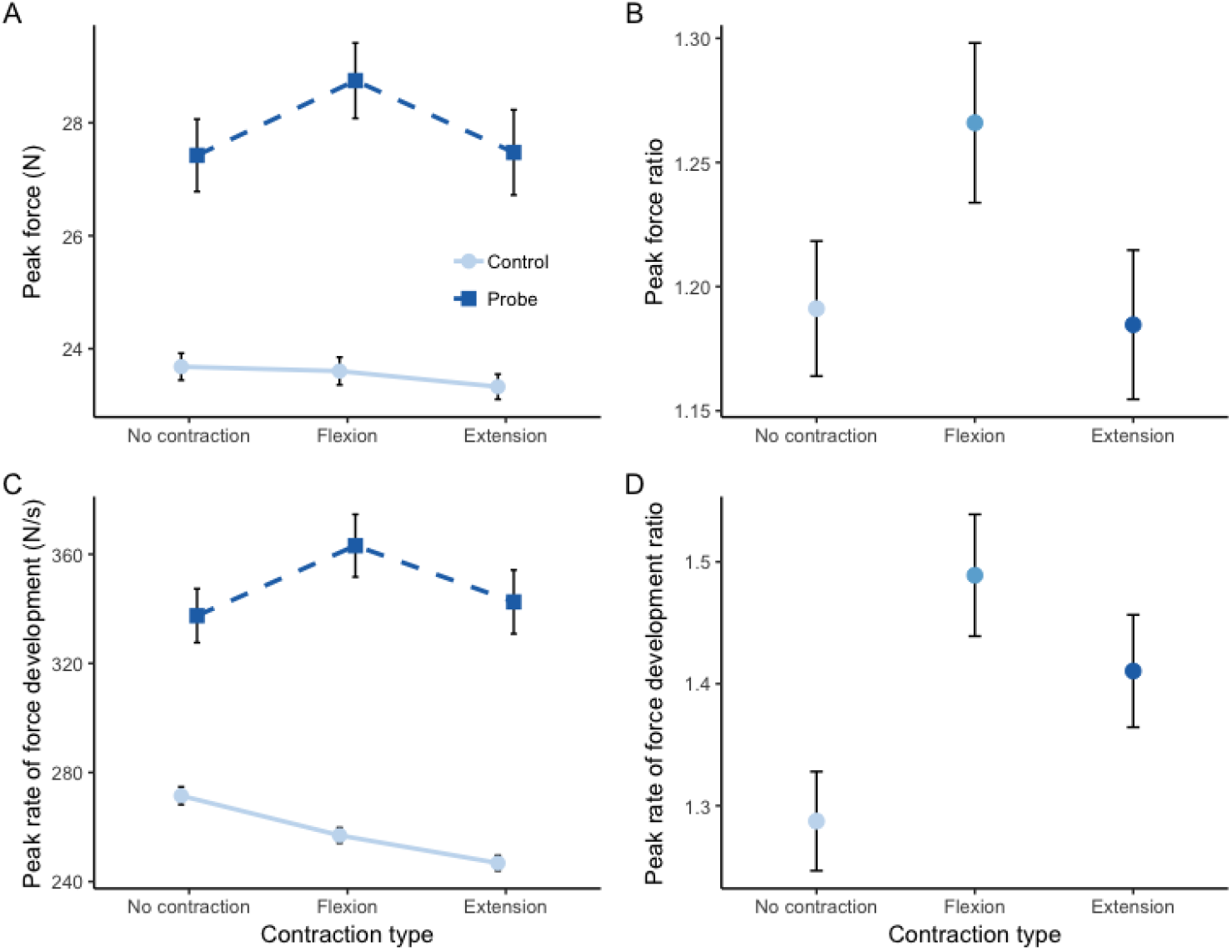
A). Mean peak force for control and probe trials for each contraction type. B). Mean peak force ratios for each contraction type. C). Mean peak rate of force development over control and probe trials at each contraction type. D). Mean peak rate of force development ratios for each contraction type. Error bars represent standard error of the mean.

## 8.0 Discussion

Intense sounds have paradoxical effects on the motor system, depending on the evolving state of the central nervous system during preparation. During maintenance of a stable low-force muscle contraction, a LAS has inhibitory effects on the corticospinal tract (Fisher et al., 2004; Furubayashi et al., 2000; Kuhn et al., 2004). In contrast, during preparation for a discrete movement, a LAS has an excitatory effect on the corticospinal pathway (Marinovic, Tresilian, et al., 2014). Facilitation of movement via intense sound (the StartReact effect) is observed when a LAS is delivered whilst the central nervous system is in a high state of preparation (close to movement initiation time ~ 200 ms). As such, this excitatory effect of sound during preparation may provide a neurophysiological means by which motor performance can be enhanced in the StartReact effect (Marinovic, Tresilian, et al., 2014; Marinovic & Tresilian, 2016). Observations of paradoxical effects of sound on corticospinal excitability which are contingent on the motor system’s state of preparation raise the question of whether combined muscle contraction and motor preparation enhance, or diminish, the StartReact effect. Therefore, here we investigated how the combined maintenance of a muscle contraction during preparation for action impacts the facilitation of motor output induced by a LAS. Raw RTs of movements executed in the bilateral task of experiment one provided no evidence that preparatory contractions of different force levels impact the degree to which a LAS can shorten RT. However, analysis of the entire RT distribution using CDFs indicated some overall benefit of a 10% MVC preparatory contraction on RT, and an overall delay of RT when a 20% of MVC contraction was maintained during preparation. Consideration should also be given to a potential RT floor effect which may have resulted in RTs being already close to the limits of the central nervous system and therefore limiting the facilitation of actions by the LAS in terms of their initiation.

In experiment two, our CDF analysis indicated that sustained flexion contractions at 10% of MVC, which produced the most benefit on RT in the bilateral task of experiment one, resulted in a delay of movement initiation across the RT spectrum when performed in the unilateral task prior to initiation of the ballistic movement. A similar delay was also produced by the sustained extension contraction during the unilateral task. Attention has previously been shown to modulate intracortical inhibitory circuits of M1 (Bell et al., 2018; Binkofski et al., 2002; Kuhn et al., 2017), with external focus of attention, as opposed to an internal one, increasing short-interval intracortical inhibition (SICI) during sustained contraction (Kuhn et al., 2017). The authors suggest that this serves to regulate the amount of M1 outflow and subsequently increase the time taken for muscle fatigue to occur. This may be applicable to our data, as participants were provided with visual feedback regarding their hand position so that the correct amount of force would be exerted during the sustained contraction. However, we removed this visual feedback prior to LAS presentation and movement onset. Therefore, preparation of the ballistic response may have produced a similar modulation of SICI – either through a shift of attention toward the impending IS presentation, or by the process of motor preparation itself. As such, these attentional-dependent effects may have been induced within the cortical hemisphere that was engaged for the prepared response in the unilateral task but not the bilateral one. This increase of SICI within intracortical circuits might explain why this delay of RT for both contraction directions was observed during the unilateral task but not during the bilateral task.

Similar to the apparent benefit on RT that was produced by the 10% MVC contraction in the bilateral task (experiment one), the LAS provided a larger facilitatory effect on peak force and vigour when a contraction 10% of MVC was maintained contralateral to the hand engaged in preparation. Interestingly, the contralateral sustained flexion contraction during preparation in experiment three replicated this magnification of the LAS effect on peak force, however, the contralateral extension contraction did not. Rather, the LAS effects on peak force of the ballistic response were no more beneficial than the simple unilateral response when an extension contraction was maintained contralaterally. Given the flexion sustained contraction enhanced the StartReact effect but the extension one did not, this may suggest that the magnification of the StartReact by such sustained contractions can be muscle (or directionally) specific. During bilateral movements, interhemispheric inhibition has been found to be greater during isometric contraction of homologous muscles (i.e. flexion-flexion and extension-extension), whereas this inhibition is decreased during contraction of non-homologous muscles (i.e flexion-extension) (Perez et al., 2014). This decrement of interhemispheric inhibition only during asymmetrical movement appears to be incompatible with the findings we present here of an increased StartReact effect during the bilateral task when the limbs are moving congruently, but not when they are moving incongruently. However, it is difficult to directly compare these findings, given the multitude of evidence which suggests that the modulation of M1 excitability is particularly sensitive to the background state of motor circuits and the dynamics of the movements which are being executed (Carson, 1995; Chen et al., 2016; Cheney & Fetz, 1980; Dettmers et al., 1996; Marinovic et al., 2014). As such, given Perez et al. (2014) employed bilateral isometric contractions whereas we used a task engaging isometric contraction of one limb during active preparation for a ballistic response of the contralateral limb, this may contribute to the incompatibility of our findings.

These data demonstrate that the inhibitory effects on motor pathways that are induced by acoustic stimulation during the maintenance of a muscle contraction can be reversed if motor preparation coincides with certain types of contractions. The engagement of a contralateral muscle contraction may engage a wider and more distributed neural network during preparation which can subsequently be more easily recruited by the LAS and add to the accumulation of preparatory neural activity which summates to produce the final magnitude of motor output (McInnes, Corti, et al., 2020). Similar suggestions have been made to describe previous observations that the facilitation of movement triggering via the StartReact effect can vary between different movement types of the same muscle, depending on the task functionality of the movement employed (e.g. Honeycutt et al.’s (2013) finger pinch versus grip task, see Marinovic, de Rugy, et al., 2014). The finding that at least in terms of peak force, a contralateral flexion contraction increases the benefit of the LAS could be a result of the efferent connectivity of the flexor muscle. For example, it has been suggested that flexor muscles receive greater functional contributions from the corticospinal tract in comparison to extensors (Godfrey et al., 2013; Koganemaru et al., 2010; McInnes, Corti, et al., 2020; McMillan et al., 2004; Park & Li, 2013; Vallence et al., 2012), which may allow a greater facilitation of force due to the correspondence of force generation with primary motor cortex (M1) activity (Ashe, 1997). Alternatively, these effects may be due to the congruency of the sustained contraction with the ballistic response.

We also observed a greater benefit of the LAS on force and vigour of the ballistic response for sustained contractions at lower force levels – particularly at 10% of MVC – than for the higher force contraction (20% of MVC). The direction of this effect is opposite to our prediction of an increase of the StartReact effect which is proportional to the strength of the contraction maintained during preparation. The use of positron emission tomography has identified that at lower force levels, there is a rapid increase of M1 activity as the amount of force produced is increased, but that the rate of this rise diminishes at higher force levels, producing a logarithmic relationship between force production and M1 activity (Dettmers et al., 1996). Single cell recordings have also suggested weak forces are primarily produced by corticomotoneurons (Maier et al., 1993), a finding which may reflect the use of weak forces in fine motor control such as precision grip (Oliveira et al., 2008; Quinn et al., 2018; Shim et al., 2007; Yu et al., 2010). Furthermore, it has been argued that the reticulospinal tract becomes increasingly important for the production of higher levels of force (Baker, 2011), given ipsilateral motor evoked potentials, which are likely mediated by the reticulospinal tract (Ziemann et al., 1999), can be more easily elicited during strong background muscle activity (Alagona et al., 2001). Therefore, the heightened use of the corticospinal tract at lower forces may have led to these force producing neurons to be more readily recruited by the LAS when engaged in a light muscle contraction during preparation, adding to the final motor output. However, this was only observed when the sustained contraction was performed contralaterally to the ballistic response, and not when it was performed ipsilaterally. Therefore, any potential interaction of both facilitatory and inhibitory effects which act during ipsilateral contraction and preparation should be considered — such as a potential modulation of SICI induced in the hemisphere that is engaged in preparation, as discussed earlier.

Finally, the sustained contractions appeared to be more beneficial to motor output when they were maintained during preparation of the contralateral limb, rather than the ipsilateral one. There are a number of neurophysiological mechanisms which may underpin this finding. For example, tonic contraction of one limb can increase activity in ipsilateral M1 (Carson et al., 2004; Kawashima et al., 1998; Liepert et al., 2001; Muellbacher et al., 2000), which may be mediated by interhemispheric modulations of excitability via the corpus callosum (Carson et al., 2004; Di Lazzaro et al., 1999; Perez et al., 2014). This increased activity in M1 may then be recruited by the LAS when triggering a movement that is prepared in those related circuits, which subsequently adds to the final output of the response. Alternatively, engagement of the motor pathways contralateral to the side that is engaging in preparation may allow activation of ipsilateral descending pathways by the LAS which may contribute to the motor output. One such descending pathway is the cortico-reticulo-propriospinal pathway, a descending tract which has been suggested to be important in functional recovery after stroke (Bradnam et al., 2013). The ipsilateral hemisphere may also contribute to motor output through the small number of corticofugal fibres which project to ipsilateral spinal motoneurons, rather than crossing at the pyramidal decussation (Phillips & Porter, 1964). These explanations assume that the input provided by the LAS is of cortical origin. A cortical origin of the descending LAS-induced activity is supported by the fact that the descending pathways which innervate primarily contralateral muscles (i.e. the corticospinal and rubrospinal tracts) receive significant projections from the cortex (Lemon, 2008). The subcortical dorsolateral pathways, in contrast, project bilaterally (Lemon, 2008) and transmission via these pathways would likely be evident regardless of whether the task was unimanual or bimanual. Alternatively, facilitation of a bilaterally projecting pathway in the bilateral task may explain why we observed facilitatory effects induced by the LAS in the bilateral task but not the unilateral one. However, we believe this explanation seems less likely based on the multiple lines of evidence we have already discussed, suggesting a potential role of the cortex in the facilitation provided by a LAS. For example, if facilitation of a bilateral pathway (i.e. the reticulospinal tract) underlies the effects we observed, then magnification of the StartReact effect would be expected to be greatest at higher force levels due to a potentially greater involvement of reticulospinal circuits at higher force levels (Baker, 2011). It is also possible that the engagement of a muscle contraction during preparation may simply raise the level of preparatory activity to a higher state and thereby enhance the magnitude of the StartReact effect. However, we deem this to be unlikely, given the modulation of gains introduced to the ballistic response by the LAS were dependent on the force and muscle of the sustained contraction, and the effect was far more pronounced in the bilateral tasks rather than in the unilateral one. Regardless of the specific mechanisms underpinning the greater facilitation of motor output provided by the LAS in the presence of a sustained contralateral muscle contraction during preparation, this finding may have important practical implications for using the StartReact effect as a rehabilitative tool. Engagement of the contralesional side can be used to increase the benefits of acoustic stimulation and further aid in the functional recovery of movement after neurological conditions such as stroke. This is particularly promising given the ipsilateral cortex has been suggested to be capable of compensating for contralateral cortex deficits after stroke (Serrien et al., 2004; Strens et al., 2003). Furthermore, given contralateral muscle activity during preparation was shown to modulate the StartReact effect at even moderately low force levels, it may be an important consideration for researchers studying the StartReact effect to observe participants during experimental sessions to ensure they are not unknowingly activating task-irrelevant muscles.

## Notes

### Competing Interest Statement

The authors have declared no competing interest.

